# Evolution of Bacterial Gene Transfer Agents

**DOI:** 10.1101/402982

**Authors:** Rosemary J. Redfield, Shannon M. Soucy

## Abstract

Bacterial gene transfer agents (GTAs) are small virus-like particles that package DNA fragments and inject them into cells. They are encoded by gene clusters resembling defective prophages, with genes for capsid head and tail components. These gene clusters are usually assumed to be maintained by selection for the benefits of GTA-mediated recombination, but this has never been tested. We rigorously examined the potential benefits of GTA-mediated recombination, considering separately transmission of GTA-encoding genes and recombination of all chromosomal genes. In principle GTA genes could be directly maintained if GTA particles spread them to GTA^−^ cells often enough to compensate for the loss of GTA-producing cells. However careful bookkeeping showed that losses inevitably exceed gains for two reasons. First, cells must lyse to release particles to the environment. Second, GTA genes are not preferentially replicated before DNA is packaged. A simulation model was then used to search for conditions where recombination of chromosomal genes makes GTA^+^ populations fitter than GTA^−^ populations. Although the model showed that both synergistic epistasis and some modes of regulation could generate fitness benefits large enough to overcome the cost of lysis, these benefits neither allowed GTA^+^ cells to invade GTA^−^ populations, nor allowed GTA^+^ populations to resist invasion by GTA^−^ cells. Importantly, the benefits depended on highly improbable assumptions about the efficiencies of GTA production and recombination. Thus, the selective benefits that maintain GTA gene clusters over many millions of years must arise from consequences other than transfer of GTA genes or recombination of chromosomal genes.

## Introduction

Some bacteria are known to produce phage-like particles that contain short fragments of chromosomal DNA (Marrs 1974; Lang et al. 2012; Lang et al. 2017; Québatte et al. 2017; Tamarit et al. 2018). Like a phage, each particle can inject its DNA into a related cell, where the DNA may recombine with the recipient’s chromosome and perhaps change the cell’s genotype (Fig. 1). GTA particles are typically encoded by a cluster of head, tail and DNA-packaging genes strongly resembling those of known temperate phages.

**Fig. 1:**
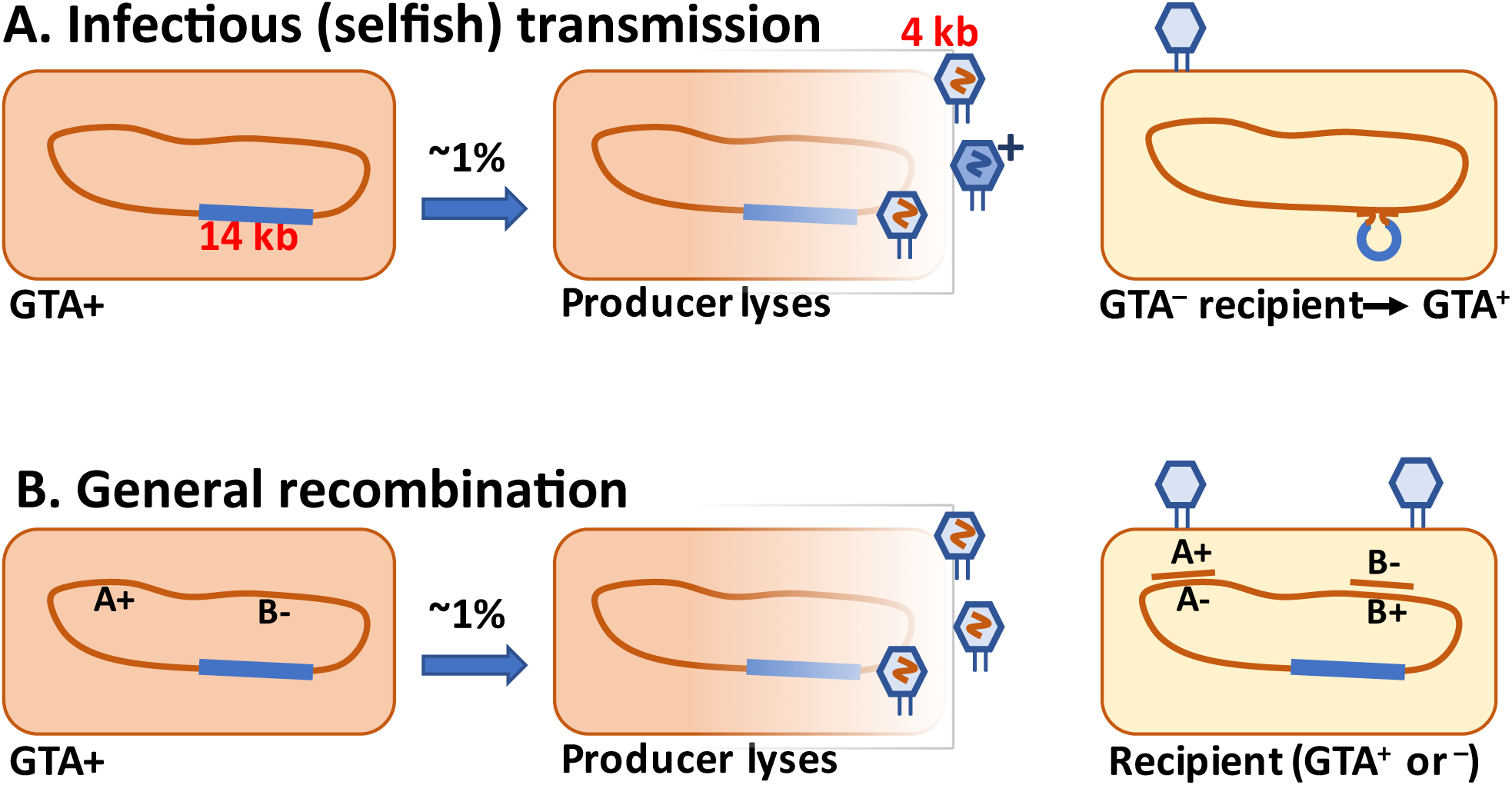
Schematic drawings of GTA-mediated recombination. **A.** Transmission of a GTA gene or cluster to a GTA^−^ cell. **B.** Transmission of chromosomal alleles to a cell with variant alleles.

The best-studied GTA system is that of *Rhodobacter capsulatus* (RcGTA; (Marrs 1974; Lang et al. 2012; Lang et al. 2017)). Genes for the RcGTA capsid (head, tail and DNA packaging) are together in a 14 kb cluster; head spikes, tail fibers, and a holin and endolysin are encoded separately (Lang et al. 2017). In lab cultures these genes are most active in stationary phase, reflecting the combined actions of at least two phosphorelays and a quorum-sensing system, all of which also contribute to regulation of other cellular processes (summarized by (Lang et al. 2017)). However, reporter-gene analyses indicate that only a small fraction of cells in a culture (0.1 - 3%) activate their GTA genes under inducing conditions (Fogg et al. 2012). Each 30 nm-diameter RcGTA particle contains an approximately 4.4 kb fragment of chromosomal DNA, with all regions of the chromosome equally represented (Hynes et al. 2012). Burst sizes are not known, but consideration of the *R. capsulatus* genome size (3872 kb), the DNA capacity of the particles, and the maximum titer of particles transducing a specific marker (about 2×10^6^/ml (Solioz et al. 1975)) suggests that culture supernatants can contain at least 2 × 10^9^ GTA particles/ml.

Persistence of GTA genes is usually attributed to selection for the chromosomal recombination they cause (hence their name) (Solioz et al. 1975; Koonin 2016; Québatte et al. 2017; Westbye et al. 2017). However, this explanation should be treated with caution for several reasons. First, although extensive investigations of eukaryotic sexual reproduction have identified some conditions where random recombination of chromosomal genes is beneficial, such conditions are much less ubiquitous than generally thought (Burt 2000; Altenberg et al. 2017). Second, GTA-mediated recombination is likely to be quite inefficient since the DNA-containing particles are released into the environment, not directly passed to recipient cells. Third, the particles are released by lysis of the producer cells; this strong fitness cost of GTA production can only be overcome by a strong benefit to the GTA genes themselves or to the recipient cells.

Below we first evaluate whether GTA-encoding genes can use GTA particles to spread themselves to GTA^−^ cells at a high enough frequency to compensate for the death of GTA-producing cells. Simple algebra indicates that this is not possible under any reasonable assumptions, because no plausible efficiency of transmission can overcome the associated loss of GTA-producing cells.

The major part of this paper then considers whether a GTA gene cluster can be maintained by benefits arising from recombination of other chromosomal genes. We focus on ways that recombination can reduce the burden of deleterious mutations, rather than increasing the benefits of beneficial mutations, since the former are much more likely to provide the recurrent advantage that would be needed to maintain GTA genes (Kondrashov 1982; Kondrashov 1988). We apply a common population-genetics model, where synergistic epistatic interactions between mutations in infinitely large populations skew the distribution of mutations, creating unfavourable conditions that recombination can remedy (Redfield 1988). Although real populations are not infinite and real genes do not have consistently synergistic epistasis, this model provides a useful benchmark to evaluate whether benefits of GTA-mediated recombination can compensate for the lysis of GTA-producing cells. We then extend the model to populations where cells’ fitness regulates their ability to produce and receive GTAs, mimicking the reported modes of GTA regulation by physiological signals. Although both epistasis and regulation models identify some conditions where GTA^+^ populations are fitter than non-recombining populations, these benefits depend on the highly improbable assumption that all steps proceed with 100% efficiency. Finally, we examine whether recombination benefits (arising from a combination of epistasis and regulation) can prevent GTA^−^ mutants from invading a GTA^+^ population, or allow GTA^+^ cells to invade a GTA^−^ population.

## RESULTS

We initially focused on two models where DNA transfer by GTA particles directly benefits the GTA genes responsible for particle production, by transferring either the whole GTA cluster or single GTA-gene alleles. Because their benefit to GTA genes is direct, such ‘selfish’ explanations have the potential to give much stronger selection than could arise from indirect benefits of general chromosomal recombination.

### Selfish transfer version 1: Transfer of a GTA gene cluster

Under this hypothesis, the GTA gene cluster persists in populations because some GTA particles (‘G^+^ particles’) package this DNA and transfer it into GTA^−^ cells, generating enough new GTA^+^ cells to compensate for the deaths of producer cells and occasional mutation or deletion of the gene cluster (see the schematic in Fig. 1A). One obvious problem with this hypothesis is that most species’ GTA gene clusters are too long to be transferred intact in the GTA particles they encode. We initially hypothesized that the GTA capsid proteins might occasionally assemble into anomalously large particles that could contain the complete GTA cluster plus enough flanking chromosomal DNA to facilitate homologous recombination. However the following simple model makes it clear that the GTA cluster cannot increase its frequency by GTA-mediated transfer even if the length constraint is removed.

The model’s simplifying assumptions were all chosen to favour transmission of GTA genes, so that its conclusions would overestimate rather than underestimate the benefits of GTA production. First, it assumes that only one gene is needed for production of GTA particles, and that the DNA capacity of these particles is substantially larger than this GTA^+^ gene. When this gene is expressed, all of the cell’s DNA is assumed to be packaged into GTA particles and released into the environment by cell lysis. The model also assumes that all other cells in the population lack the GTA^+^ gene, but can accept and recombine DNA from GTA particles, as if the GTA^+^ cell has just entered a naive GTA^−^ population. It further maximizes efficiency of GTA transmission by assuming that all GTA particles survive in the environment, successfully attach to recipient cells, and inject their DNA, and that all injected DNA strands replace their chromosomal homologs.

In this maximally favourable model, what conditions will lead to an increase in the number of GTA^+^ cells? Since producer cells die, and all G^+^ particles are assumed to attach to GTA^−^ cells and convert them to GTA^+^, the critical value is the number of G^+^ particles produced by each GTA-producing cell; GTA^+^ cells can increase only if this is greater than one. However, in cells with unreplicated genomes this value cannot be greater than one, because each GTA^+^ cell’s genome will contain only a single copy of the GTA genes. The GTA gene-cluster is not amplified before GTA production (Hynes et al. 2012), and the GTA genes do not encode any self-replication mechanism. Thus, under our assumption of perfect efficiency, one GTA^+^ cell must die to produce each G^+^ particle and at most one new GTA^+^ cell can be created by it. This would be just sufficient for GTA^+^ maintenance if no other factors removed GTA^+^ cells from the population, but it would not compensate for random GTA mutations and deletions and the occasional extinction of GTA^+^ lineages.

All changes that make the model more realistic would further reduce the ratio of new GTA^+^ cells to lysed ones. If the G^+^ gene is assumed to already be common in the population, many G^+^ particles will wastefully inject their GTA^+^ DNA into cells that are already GTA^+^. Considering a typical long GTA gene cluster instead of a single GTA^+^ gene would require that some or all GTA particles be large enough to contain a full-length cluster plus enough flanking DNA to promote homologous recombination with the recipient chromosome, but such large particles have never been observed. Relaxing the assumptions of perfect packaging and transduction efficiency would further reduce the odds, perhaps by several orders of magnitude. Even if it were the case that the genome was replicated without cell division prior to GTA production the upper limit would be two G^+^ particles. This over-replication would only increase actual GTA production if the expression of the GTA structural genes were also proportionately increased, and it is unlikely to compensate for the inability to package the entire GTA genome, and the G+ particles being injected into GTA+ cells.

### Selfish transfer version 2: Preferential transfer of functional GTA alleles into GTA^−^ mutants

The critical limitation of the simplified model above also applies to less restrictive situations where all the cells in the population contain a GTA gene cluster, but some cells’ clusters contain defective mutant alleles. These mutant alleles can be restored by GTA-mediated transduction with DNA fragments containing GTA^+^ alleles from GTA producer cells. The converse (replacement of functional alleles by defective ones) would not occur since cells with defective alleles would not produce GTA particles. This asymmetry favours gain of GTA^+^ over loss, and is the inverse of the ‘Sex with poor relations’ situation that arises for genes conferring natural competence (Redfield et al. 1997). However, it faces the same limitation as transfer of complete GTA genes, since each GTA^+^ cell can only produce one particle containing any particular restoring allele. Thus, the frequency of this transduction will necessarily be less than the loss of GTA-producer cells by lysis. Efficiency will be particularly low because this situation is most likely to arise by mutation of GTA^+^ alleles in a primarily GTA^+^ population, causing most of the G^+^ particles to waste their DNA on GTA^+^ cells.

The analysis of these two model versions shows that it is not enough for a GTA system to be potentially self-transmissible, able to fit the full gene set into a single GTA particle. The fundamental factor differentiating the GTA situation from other infectious-transfer models is that the GTA genes have no autonomous replication mechanism. They are replicated only along with the rest of the cell’s chromosome, and thus can only be maintained against random mutation and deletion if they increase the fitness of the genomes they are in. The rest of this work investigates ways that GTA-mediated recombination of non-GTA genes (Fig. 1B) can increase fitness.

### Modeling the benefits of general recombination

Figure 1B schematically shows GTA-mediated recombination of chromosomal alleles. Although recombination can create new combinations of new beneficial mutations, such mutations arise rarely and thus are unable to provide the consistent advantages needed to maintain the genes causing recombination (Kondrashov 1982). More reliable fitness effects can be created by recombination of deleterious alleles, which are continually arising by mutation even in the best-adapted genomes (Robert et al. 2018). Development of our deleterious-mutation model was guided by the deterministic transformation model of Redfield 1988. It does not explicitly address the evolution of genes causing GTA, but instead compares the mean fitnesses of pure populations that are either GTA^+^ or GTA^−^. In such populations benefits that arise from recombination of chromosomal alleles can increase the population’s mean fitness above the equilibrium set by the mutation rate.

The main steps in the model are labelled in Fig. 2, key parameters are listed in Supplementary Table I, and details are diagramed in Supplementary Figure A and explained in its legend. The model considers a constant-size population large enough that stochastic processes such as genetic drift can be ignored. Cells begin with no mutations, and accumulate an average of 1.0 deleterious mutations per generation; this is substantially higher than real mutation rates (Lynch et al. 2016), increasing the potential benefits of recombination. All mutations have the same fitness cost ***s*** (***s*** = 0.1); this allows all cells with the same number of mutations to be considered together (individual cells and mutations are not tracked). A cell’s fitness depends on how many mutations it has, and is described by a multiplicative fitness function ***w*** = (1 - ***s***)**^*i^α*^**, where ***i*** is the number of mutations in the cell’s genome and **α** is an epistasis superscript coefficient that specifies how interactions between mutations affect fitness. The fitness functions used are shown in Fig. 3.

**Fig. 2:**
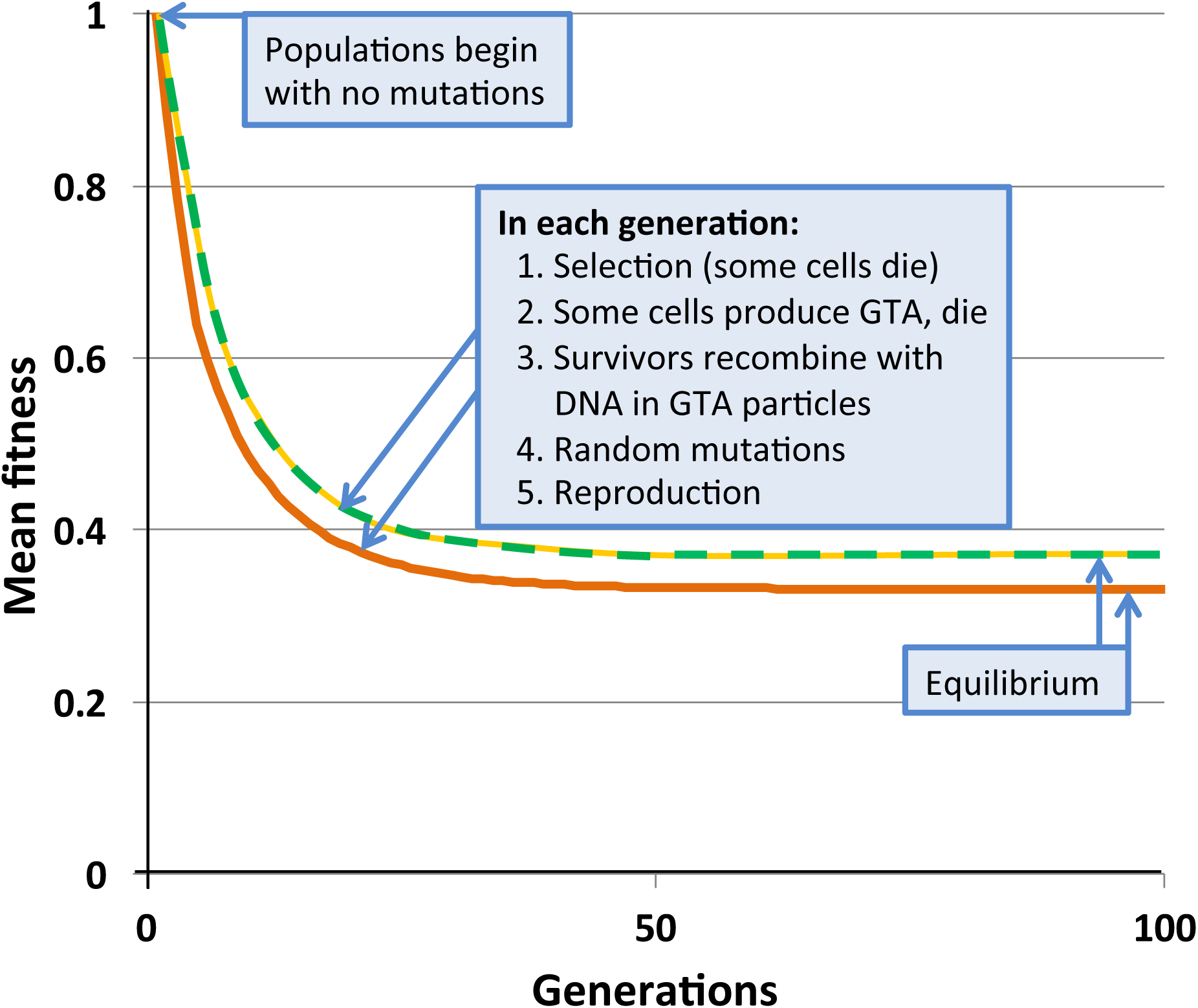
Changes in mean population fitness under simple multiplicative fitness (no epistasis). Green dashed line: no GTA production; equilibrium fitness = 0.3678. Yellow dashed line: GTA production by 10% of the population without cell lysis; equilibrium fitness = 0.3678. Orange line: GTA production by 10% of the population with cell lysis; equilibrium fitness = 0.3311.

**Fig. 3:**
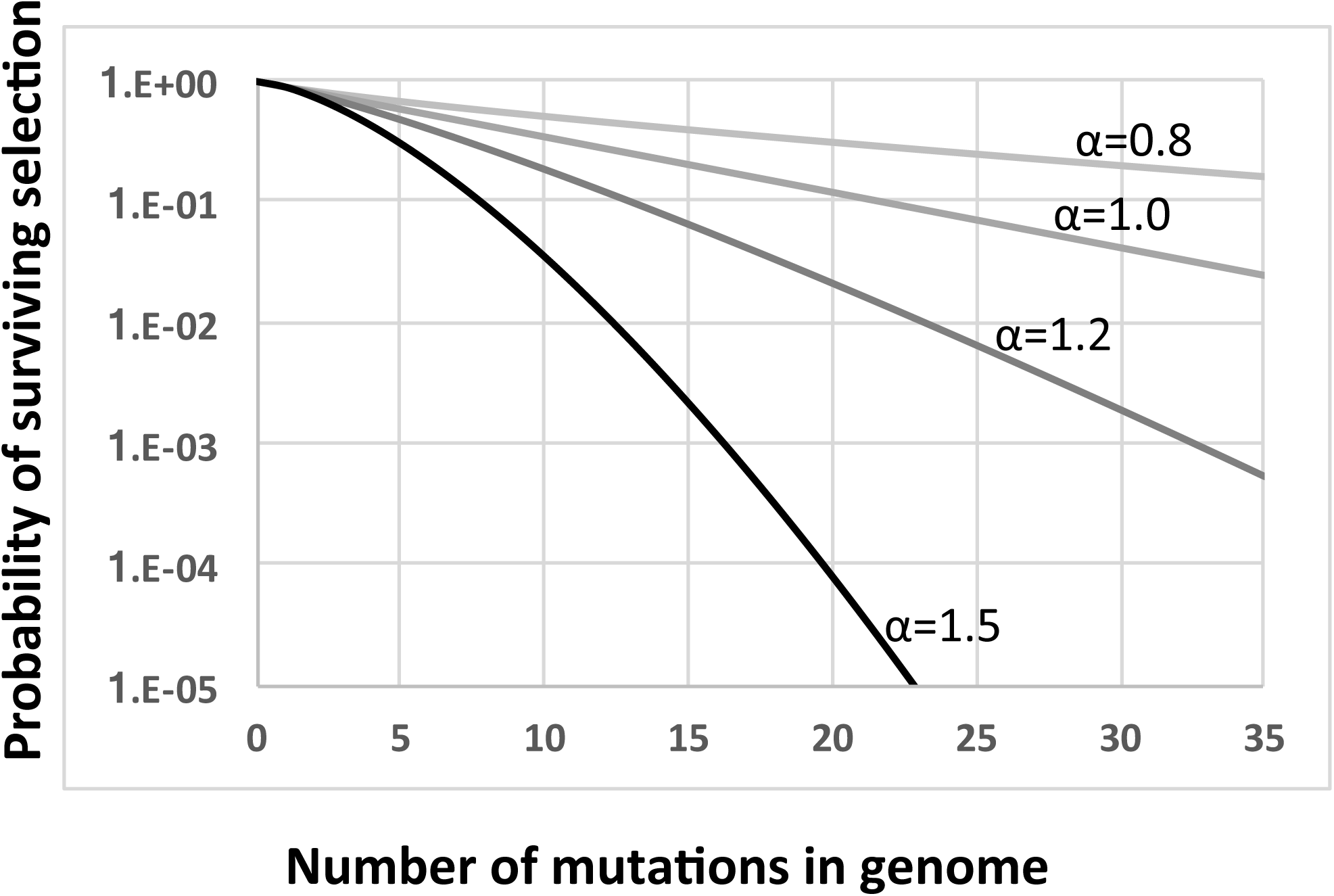
Fitness functions used in this work. All functions are based on ***w*** = ((***i*** - ***s***)**^*i*^**)**^*α*^**. Light grey line: ***α*** = 0.8. Medium grey line: ***α*** = 1.0. Dark grey line: ***α*** = 1.2. Black line: ***α*** = 1.5.

Figure 2 shows the results of example runs of the simulation using a simple multiplicative fitness function (**α** = 1). Mutations accumulate over many generations until the population reaches a mutation-selection equilibrium; at this state the number of new mutations introduced in each generation equals the number of mutations removed by selective deaths. In a non-recombining population, the mean fitness at equilibrium has been shown mathematically to equal the fraction of the population that receives no new mutations in one generation (Haldane 1937; Crow 1970). When *μ* = 1.0, this value is 0.3678, a prediction exactly matched by the simulated non-recombining population indicated by the green dashed line in Fig. 2.

### GTA production and recombination

In each model generation, some cells activate their GTA genes (with probability ***pGTA***), package their entire genome into GTA particles, and lyse. The distribution of mutations in the resulting GTA particles is calculated with a matrix giving the binomial probabilities of producing particles with ***j*** mutations from a genome with ***i*** mutations. For most tests ***pGTA*** was varied from 0 to 0.15; setting ***pGTA*** = 0 provided a no-recombination control. For comparison 0.1-3% of *R. capsulatus* cells produce RcGTA (Fogg et al. 2012), and 6-14% of *B. henselae* cells produce BaGTA (Québatte et al. 2017).

Since our goal was to identify conditions where recombination increases fitness, the model maximized the simulated recombination efficiency by assuming that surviving cells take up all the DNA in the GTA particles. This was modeled by setting the fraction of each cell’s genome replaced by recombination to also equal ***pGTA***. The binomial matrix described above was also used to calculate how recombination with DNA in GTA particles changed the mutation distribution of the surviving cells.

When ***α*** = 1 (medium grey line in Fig. 3) there is no epistasis, and each new mutation reduces fitness by the same factor, regardless of how many other mutations are in the genome. It is well established that, in the absence of epistasis, recombination in such models will not change the equilibrium fitness of a population, because the distribution of mutations remains random (Smith 1968; Redfield 1988). The yellow dashed line in Fig. 2 and the blue dots and line in Fig. 4 show control no-epistasis (***α*** = 1) simulations where recombination occurred but lysis did not (as if cells magically released GTA particles to the environment without cell lysis). The blue dashed line in Fig. 4B shows that, as expected, all values of ***pGTA*** gave the same equilibrium mean fitness as populations with no recombination. The orange lines in Fig. 2 and 4B show the more realistic situation where all cells that have produced GTA are removed from the population. With this lysis-simulating step included, the mean fitness for pGTA = 0.1 falls to a lower equilibrium (Fig. 2, orange line), and the equilibrium mean fitnesses for a range of pGTA values fall linearly as GTA production and recombination increase, due to the increasing fraction of cells that lyse.

**Fig. 4:**
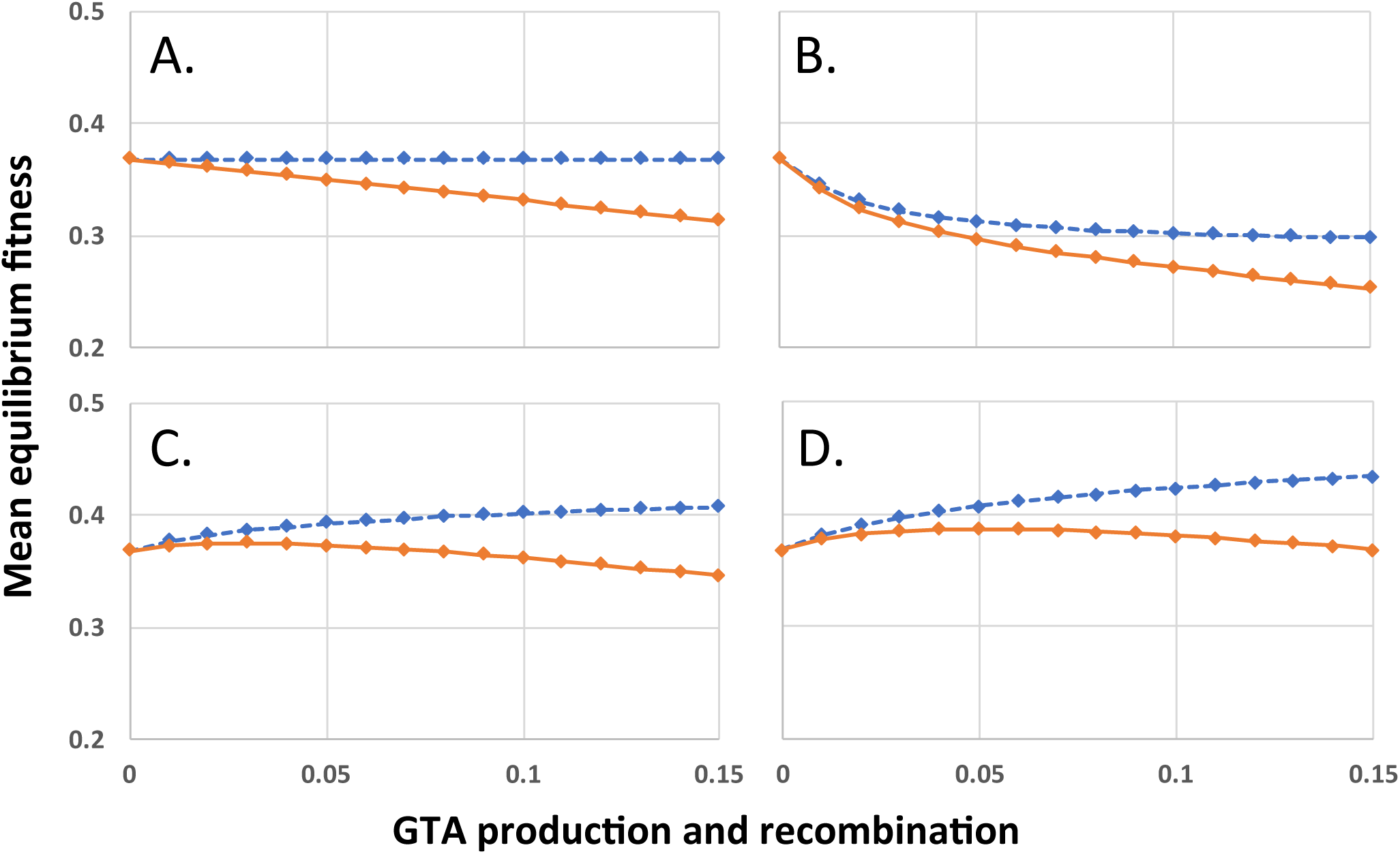
Mean equilibrium fitnesses of populations as a function of GTA production and recombination for different fitness functions. X-axis: ***pGTA***, the fraction of cells producing GTA and the fraction of genome recombined in surviving cells. **A**. ***α*** = 0.8; **B. *α*** = 1.0; **C. *α*** = 1.2; **D. *α*** = 1.5. Blue: GTA production without cell lysis; orange: GTA production with lysis.

### Recombination benefits arising from epistasis

Mathematical work has also shown that, when epistatic interactions cause the effect of a new mutation to depend on how many other mutations are already present, recombination will increase or decrease equilibrium mean fitness, depending on the direction of the epistasis effect (Eshel and Feldman 1970). Values of ***α*** less than 1 give conditions where the impact of each additional mutation is reduced in proportion to the number of other mutations already present (light grey line in Fig. 3). With this ‘antagonistic’ epistasis, recombination will reduce equilibrium mean fitness below that of a non-recombining population.

Selection with high epistasis improves the equilibrium fitness of the population because it imposes a higher per-mutation cost on cells with heavy mutation loads. This shifts the distribution of mutations in the population so it is skewed towards lower mutation loads (no longer random). In the absence of recombination this reduces the average number of mutations per cell but this reduction does not change the mean equilibrium fitness because the skew makes selection less efficient. Recombination removes the skew, randomizing the distribution around the new lower mean load and increasing the proportion of cells with high numbers of mutations. This allows selection to more efficiently remove mutations from the population (one cell death removes many mutations), resulting in a higher population fitness at equilibrium. Supp. Fig. B. shows the effects of different levels of epistasis on equilibrium mutation distributions; outcomes are shown in Fig. 4.

The effect of antagonistic epistasis (***α*** = 0.8) is shown in Fig. 4A. Increasing GTA production reduces the population’s equilibrium fitness even in the absence of lysis (blue dashed line), and reduces it more strongly when lysis occurs (orange line). Fig. 4C and 4D show results for ***α*** = 1.2 and 1.5 respectively. In both cases the equilibrium mean fitness rises with increasing GTA production, but the rapid increase for low values of ***pGTA*** slows as recombination increases, and is eventually overcome by the linearly increasing cost of cell lysis (compare blue and orange lines).

This result shows that benefits of recombination can, in ideal cases, overcome the cost of GTA production. Before examining the effects of relaxing the model’s assumptions, we considered another way in which benefits might arise.

### Recombination benefits arising from regulation of GTA production and uptake

Where GTA production in real cells has been investigated, it has been found to be subject to various controls. In *R. capsulatus*, RcGTA levels are highest at the onset of stationary phase, and production is increased by nutrient stress (amino acid depletion and the stringent response). Both the RcGTA cluster and the capsule polysaccharide recognized by the particles are regulated by *ctrA* (a regulator of regulators) and by a quorum-sensing system (reviewed by Lang et al. 2017). This regulation has been interpreted as a mechanism to maximize expression at high cell densities and under stressful conditions. In contrast, production and uptake of the *Bartonella* GTA is highest in rapidly growing cells, which has been interpreted as a mechanism to maximize transmission of beneficial alleles (Québatte et al. 2017).

Could a population’s equilibrium mean fitness be improved by restricting GTA production and/or recombination to either low-fitness or high-fitness subpopulations? To test this, we introduced fitness-based criteria for participation in these steps, with low-fitness induction corresponding to the *R. capsulatus* pattern and high-fitness induction corresponding to the *Bartonella* pattern. A pair of logistic functions were used to specified how GTA production or recombination depended on mutation load (illustrated in Fig. 5). The high-fitness function (‘high-log’) identifies cells whose genomes are well-matched to their environment, with few or no deleterious mutations; the low-fitness function (‘low-log’) identifies cells whose genomes are poorly matched to their environment, with many deleterious mutations. Because each function’s shape depends on the mean number of mutations in the population, its values were calculated anew for each generation. For GTA production, this fitness regulation was superimposed on the probability of activating the GTA genes (*i.e.* the regulation determined the subpopulation to which the ***pGTA*** probability would then be applied). We determined the equilibrium fitnesses resulting from all 9 combinations of these functions in the absence of epistasis (Fig. 6).

**Fig. 5:**
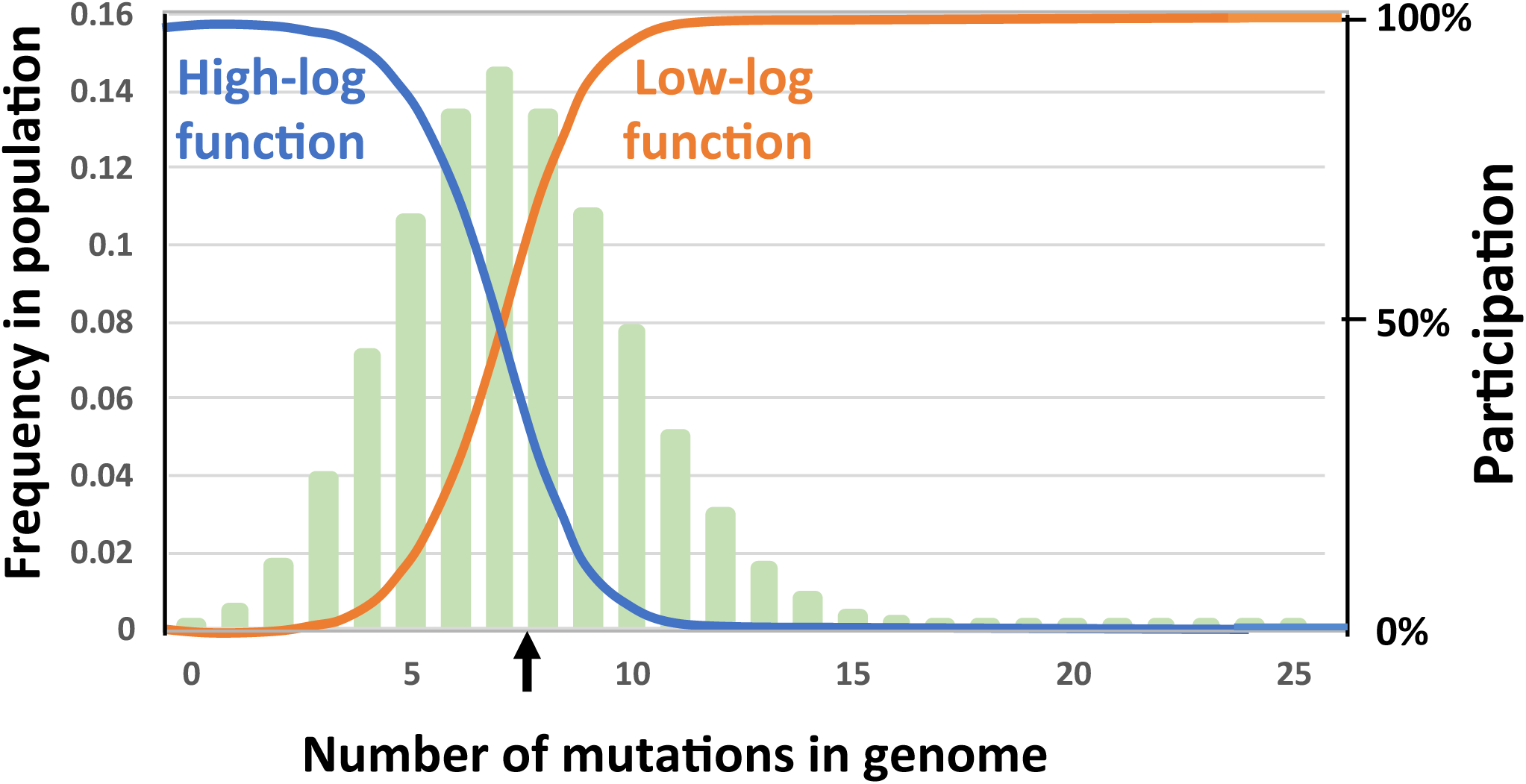
Regulatory functions that limit participation in GTA-production and/or GTA-recombination. Functions are indicated by blue and orange lines superimposed on a typical equilibrium mutation distribution (pale green bars). Left Y axis: The mutation distribution (frequency of each mutation class in the population). Right Y axis: The regulation function (relative probability of producing GTA or recombining). The black vertical arrow indicates the mean mutation number in the population. Probability of participation is a logistic function of the mean mutation load and the maximum number of mutations being considered (25 in this example). Blue: Participation mainly by high-fitness cells; orange: participation mainly by low-fitness cells.

**Fig. 6:**
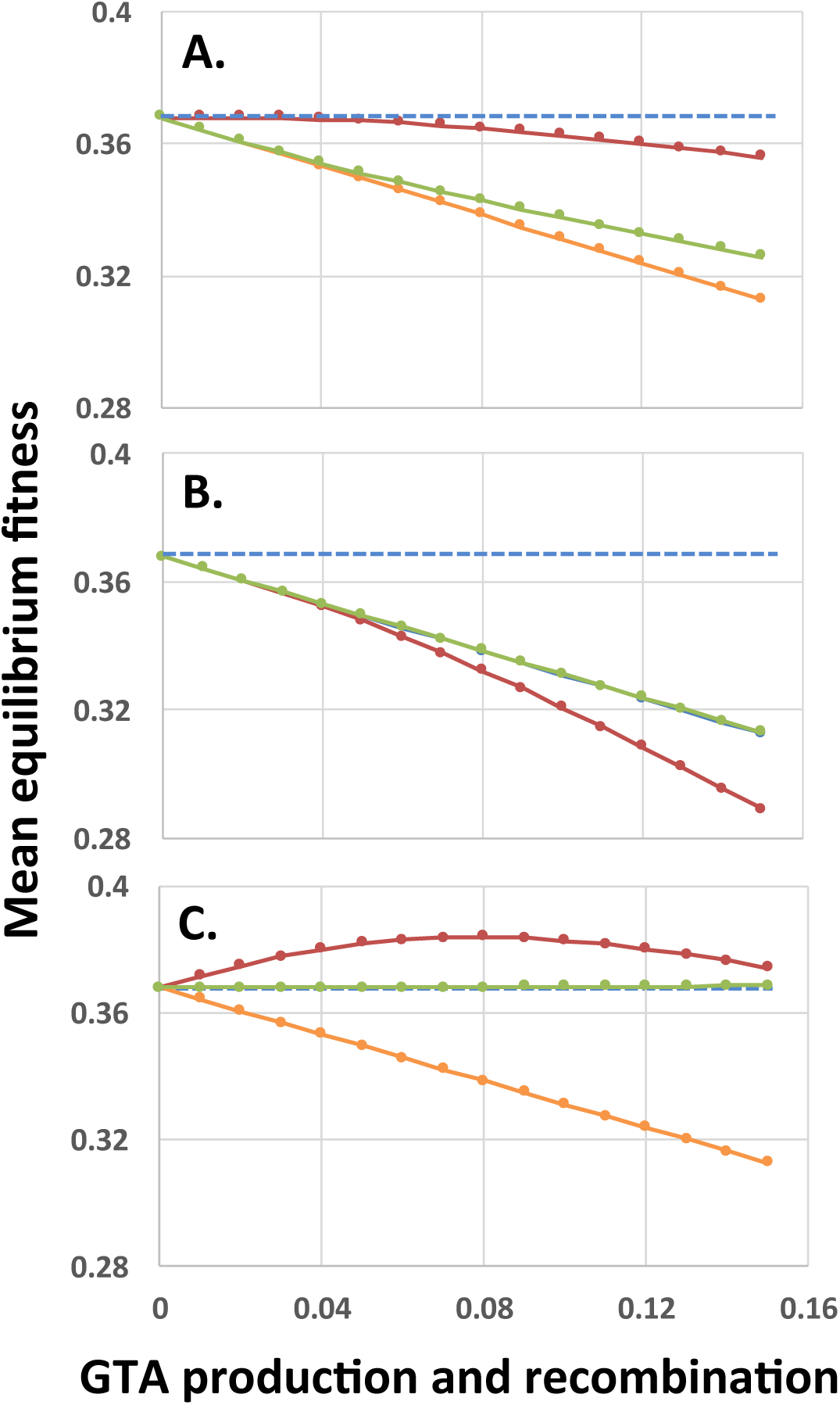
Fitness effects of regulating participation in GTA production and/or GTA uptake and recombination. GTA-producing cells lyse, and ***α*** = 1.0. **A.** Effects of regulating only GTA production. **B.** Effects of regulating only GTA uptake and recombination. **C.** Effects of coordinately regulating both GTA production and GTA uptake and recombination. Red lines: regulation by the High-log function in Fig. 5; Orange lines: no regulation, all cells participate (hidden by green line in panel B); Green lines: regulation by the Low-log function in Fig. 5. Blue dashed lines: fitness in the absence of GTA production.

In principle, regulation can alter several factors that impact fitness. Factors 1 and 2 are the quality of the DNA in the GTA fragments and in the cells that recombine with this DNA – for example, DNA from high-fitness cells will more often increase the fitness of low-fitness cells. Factor 3 is the fitness distribution of the cells that lyse – loss of high-fitness cells will reduce population fitness more than loss of low-fitness cells. Factor 4 is the effect of recombination on population fitness – because regulation skews the mutation distribution in the DNA sources and the recombining population, it allows recombination to change the distribution of mutations even in the absence of epistasis.

Fig. 6 shows the effects of regulating GTA production, GTA recombination, or both (Panels A, B or C). In each panel the blue dashed line is fitness in the absence of GTA production, and the orange line is the same data as Fig. 4B (no epistasis and no regulation; all cells participate but there is no benefit of recombination). In Fig. 6A, only GTA production is regulated. Restricting GTA production to high-fitness cells (red line) increased fitness just sufficiently to overcome the cost of lysis when pGTA is low, but not when pGTA is high. Importantly, in the absence of lysis this increase of fitness was linear with increasing GTA production (Supp Fig. C, panel B), suggesting that it was a direct consequence of the higher DNA quality (Factor 1), not an indirect effect of recombination (Factor 4). Restricting GTA production to low-fitness cells (green line) gave a much smaller benefit, though still slightly better than with no regulation. Comparison to the no-lysis control (Supp Fig. C, panel H) show that under this regulation the cost of lysis became very small, which suggests that the benefits may arise mainly from restricting lysis to the unfit cells that would otherwise die from selection.

Panel B of Fig. 6 shows the effects of regulating only recombination. When mainly high-fitness cells took up DNA and recombined, but GTA particles came from all cells (red line), equilibrium fitnesses were worse than with no regulation or no recombination. Fitnesses decreased even when GTA particles were produced without lysis (Supp Fig. C, panel D); this may partly be because fit cells are replacing their good genes with mutated genes. When mainly low-fitness cells took up DNA and recombined (green line) fitnesses were the same as in the unregulated condition (the orange line is behind the green line), perhaps because the recombining low fitness cells are already destined to die by selection.

What about when both GTA production and recombination are regulated (Panel C of Fig. 6)? When GTA production and recombination are limited to high-fitness cells (red line), fitness is increased even though producer cells lyse. The curve is very similar to that seen with strong synergistic epistasis (***α*** = 1.5) in Fig. 4D, with benefits decreasing when the amount of GTA production gets too high. The green line shows the opposite regulation pattern, limiting both GTA production and recombination to low-fitness cells. Here recombination is just beneficial enough to overcome the cost of cell lysis (the blue line is behind the green line).

No benefits were seen with the two ‘contrary’ regulatory combinations (‘high-low’ and ‘low-high’). Providing high-fitness DNA to low-fitness cells (Supp. Fig. C panel A) has surprisingly little effect on population fitness in the absence of lysis, again perhaps because the low fitness cells are already doomed. In contrast, fitness is drastically reduced when only low-fitness DNA is provided to only high-fitness cells (Supp. Fig. C Panel I).

Because of their biological relevance, we investigated in more detail the effects of high-high and low-low regulation, supplementing the logistic regulation functions with linear and cutoff functions to cover a wider range of possible regulatory effects (illustrated in Supp. Fig. D). However, these did not change the conclusions drawn from the logistic function regulation. All combinations of low-low regulation of GTA production and uptake overcame the cost of lysis but caused only infinitesimal changes in equilibrium population fitness (Supp. Fig. E, panels A-I; note the very narrow range of the Y-axes), and all combinations of high-high regulation increased fitness (Supp. Fig. E, panels J-R).

The above analyses showed that both synergistic epistatic interactions and high-fitness-dependent regulation can create conditions where GTA-mediated recombination increases population fitness despite the extreme cost of producing GTA particles. Combining both kinds of benefits further increased equilibrium fitness (Fig. 7). We next tested the effects of relaxing the model’s assumptions.

**Fig. 7.**
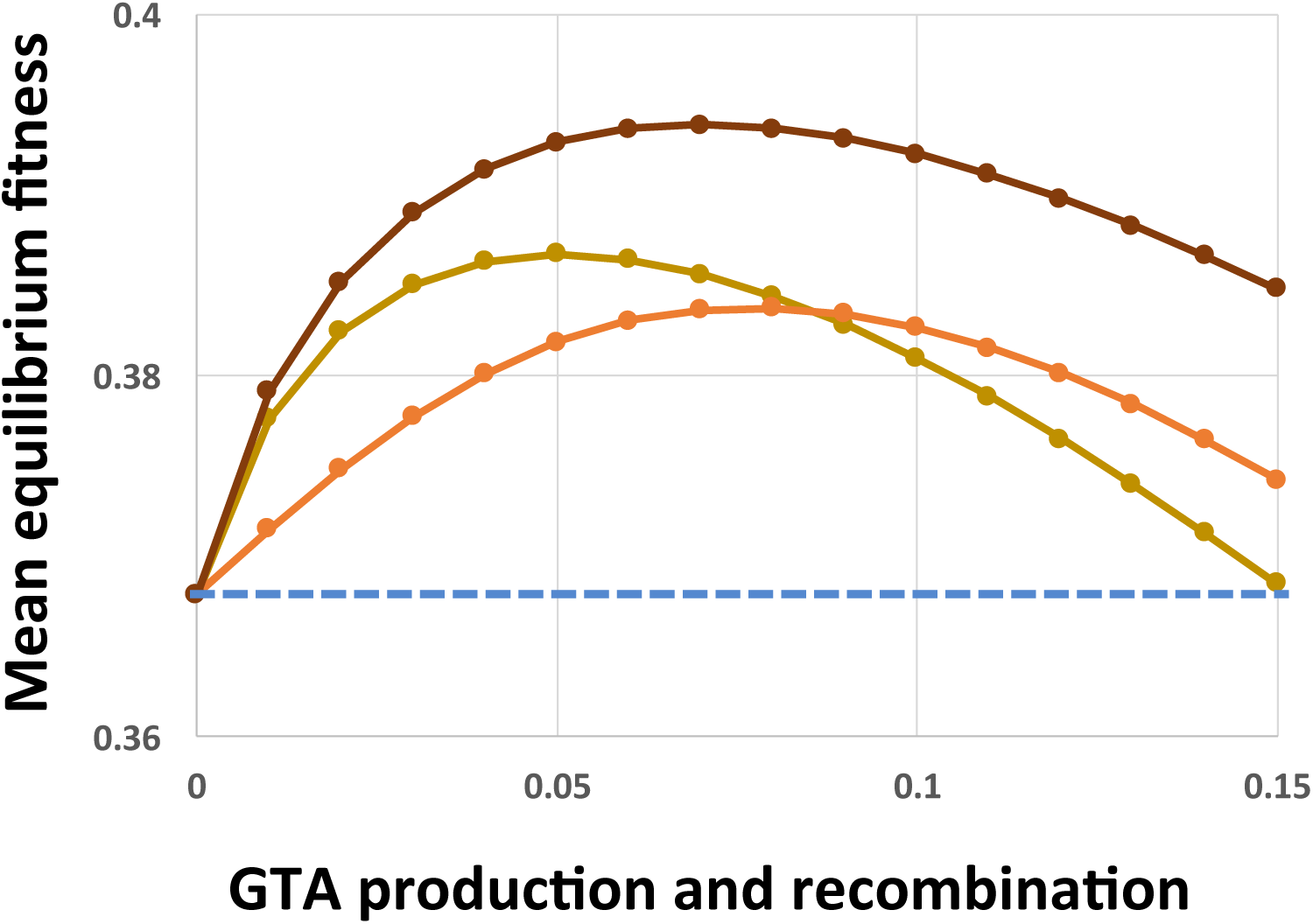
Combined epistasis and regulation benefits. Light brown line: ***α*** = 1.5, no regulation. Orange line: ***α*** = 1.0, coordinate high-cutoff regulation of both GTA production and recombination. Dark brown line: ***α*** = 1.5, coordinate high-cutoff regulation of both GTA production and recombination. Dashed blue line: no GTA production.

### Relaxing the assumptions

The recombination benefits identified above should depend on the assumptions underlying the model, since these were chosen to maximize benefits and minimize costs. We examined the impacts of three of these factors: survival of GTA particles from one generation to another, the efficiency of GTA production and recombination, and the competitive successes of GTA^+^ and GTA^−^ cells in mixed populations.

### GTA turnover

In the regulated simulations described above, any leftover GTA particles were discarded before the next generation (if, for example, more GTA particles were produced than were taken up by the recipients). Introducing a ‘turnover’ control that caused 50% of any unused GTA particles to survive to the next generation did not change the equilibrium mean fitness of any simulation, though it sometimes slightly slowed the approach to equilibrium.

### Efficiency of GTA production and recombination

The basic model assumed that GTA production and recombination have perfect efficiency, with all the chromosomal DNA of the producer cells being packaged and recombined into the recipient genomes. Much lower efficiencies will be typical of the real biological processes involved; substantial losses are very likely at production and assembly of capsid proteins, packaging of chromosomal DNA into GTA particles, survival of particles in the environment, encounters of particles with suitable recipient cells, injection and transport of DNA into the recipient cytoplasm, and eventual recombination or degradation of the DNA (see Discussion).

To capture the combined effects of these limitations, a single efficiency term ***eff*** was introduced into the model to specify the fraction of the DNA in the producing cells’ genomes that successfully recombines with recipient cells. Figure 8 shows the effects of reducing efficiency to different extents, for the combined epistasis and regulatory functions giving the highest benefits. Even 20% efficiency (red line), which is still much higher than expected for real GTA transmission (see Discussion), caused the benefits to be almost completely overwhelmed by the cost of lysis (a tiny benefit remained at the lowest amount of recombination). The cause is simple: when efficiencies are low, populations experience the full lysis costs but obtain only reduced recombination benefits.

**Fig. 8.**
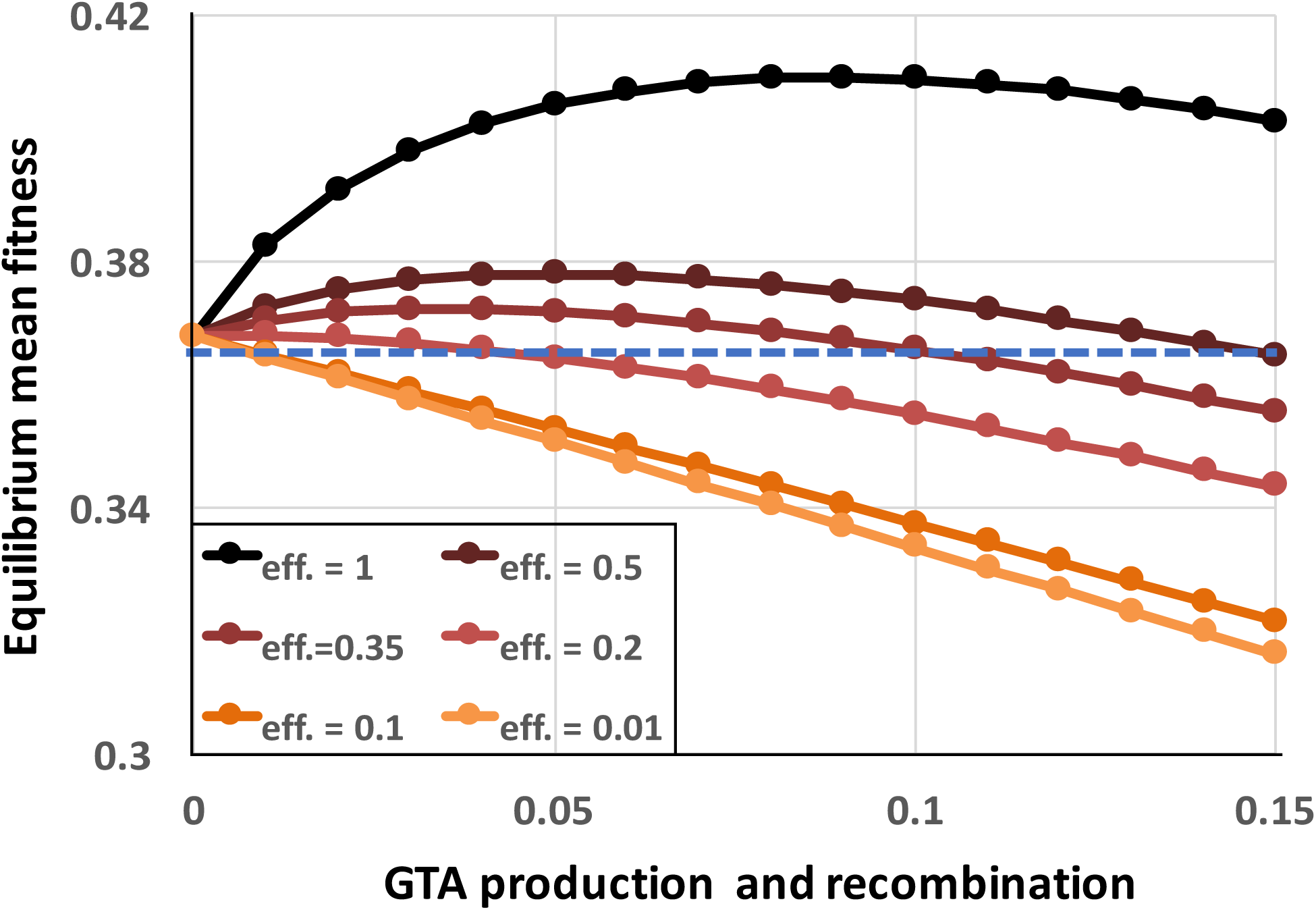
Effects of reduced GTA packaging and transduction efficiencies. *α* = 1.5, coordinated high-high regulation. Black line: *eff*. = 1.0 (as in Fig. Supp-E), dark brown: *eff*. = 0.5, dark red: *eff*. = 0.35, red: *eff*. = 0.2, dark orange: *eff*. = 0.1, light orange: *eff*. = 0.01. Blue dashed line: no GTA production.

### Competitiveness in mixed populations

The pure-population model we have considered to this point does not directly address the evolutionarily critical question of whether cells that produce GTA can outcompete non-producers. This is likely to be complex, in part because the epistasis-based recombination benefits depend on population structure, not on individual genotypes, and in part because only GTA producers experience the lysis cost but all cells able to accept DNA from GTA particles can get the recombination benefits.

To determine whether equilibrium populations of GTA^+^ cells could resist invasion by rare GTA^−^ cells, we simulated GTA^−^ and GTA^+^ subpopulations for 50 generations under the conditions of the most-beneficial GTA^+^ equilibria described above, as if we were following a small group of GTA^−^ or GTA^+^ cells in the large GTA^+^ population. Subpopulations were assumed to be so small that their activities did not affect the bulk properties of the large equilibrium population.

Fig. 9A shows the results of simulating small GTA^−^ subpopulations (dark and light green) invading a GTA^+^ population (orange) that is experiencing the combined fitness benefits of synergistic epistasis (***α*** = 1.5) and high-fitness regulation. GTA^−^ cells could invade if they took up and recombined the DNA in the GTA particles from the main population (dark green), because they did not incur the cost of lysis. If the GTA^−^ cells could not take up DNA (light green), their initial fitness advantage was gradually lost as they accumulated more mutations. The orange data points show that a control GTA^+^ subpopulation remained at constant size.

**Fig. 9:**
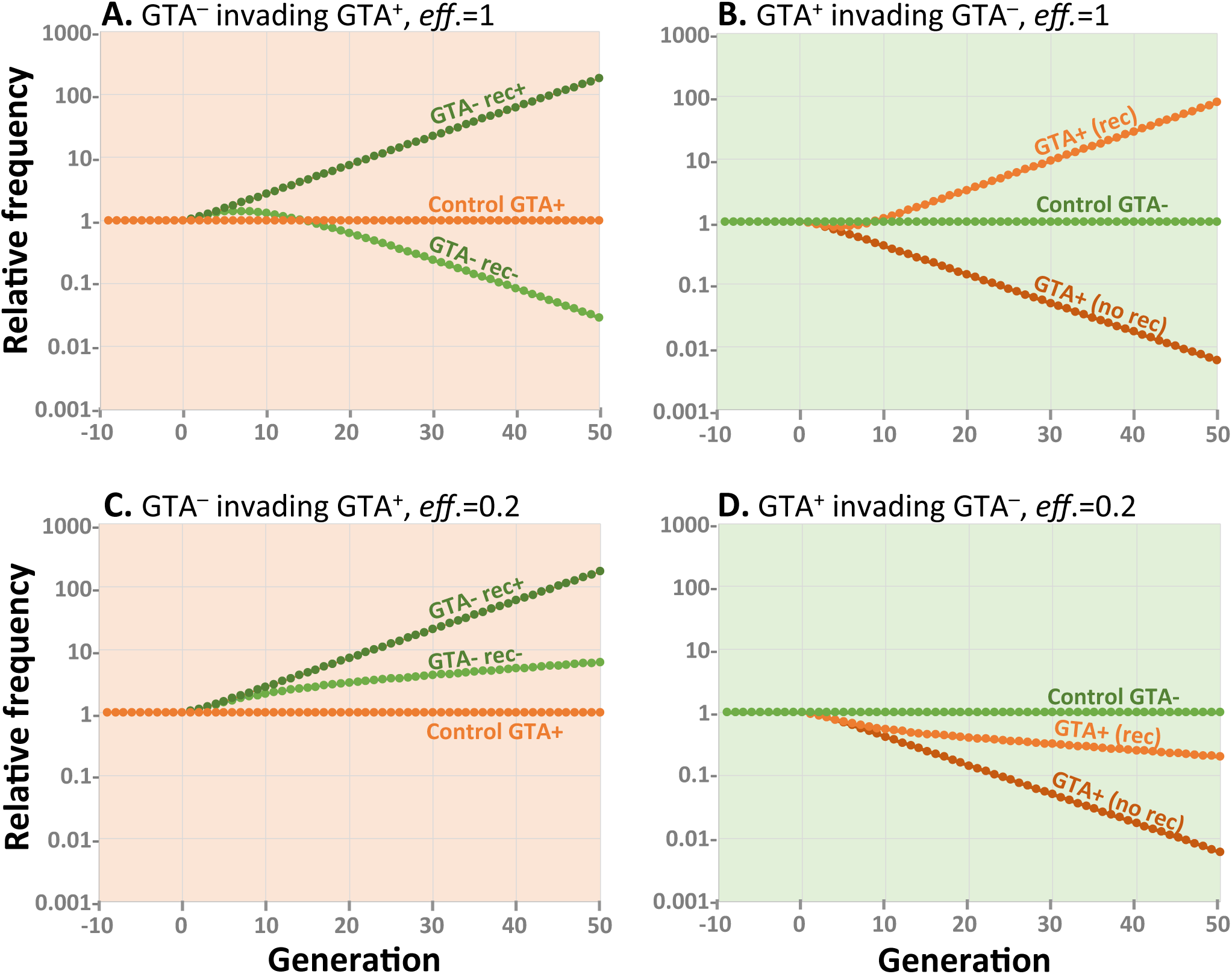
Effects of competition. All runs used ***α*** = 1.5 and high-high regulation. Note the exponential scale of the Y-axis. Panels A and B: *eff*. = 1.0. Panel A. a large GTA^+^ population being invaded by control GTA^+^ cells (orange) and by GTA^−^ cells that can (dark green) or cannot (light green) recombine. Panel B. a large GTA^−^ population being invaded by control GTA^−^ cells (green) and by GTA^+^ cells that can (light orange) or cannot (dark orange) sequester recombination. Panels C and D: Like panels A and B but with *eff*. = 0.2.

Fig. 9B shows the reverse situation, GTA^+^ subpopulations (dark and light orange) attempting to invade a large GTA^−^ population (green). GTA^+^ cells could not invade if the GTA particles they produced diffused away or were taken up by the GTA^−^ cells (dark orange). However, invasion was possible if the GTA^+^ cells could somehow sequester their GTA particles, keeping the benefits of recombination for themselves by preventing GTA-cells from recombining (light orange). The green data points show that a control GTA^+^ subpopulation remained at constant size.

Figures 9C & D show how reducing the efficiency of GTA-mediated recombination to 20% changes the competitive landscape. When the rewards of recombination are reduced relative to the cost of GTA production, GTA^−^ cells increase in a GTA^+^ population even if they cannot take up GTA DNA (Fig. 9C) and GTA^+^ cells decrease in a large GTA^−^ population even if they can sequester the benefits of recombination (Fig. 9D). As with previous simulations, the outcomes of control competitions favour GTA^+^ cells when the producer-lysis step is eliminated from the model.

Taken together, the results in Figs. 8 and 9 show that, under realistic conditions, the benefits of recombination will not be able to maintain GTA+ populations.

## DISCUSSION

Most microbiologists remain confident that horizontal gene transfer is consistently beneficial and strongly selected, despite the 50-year failure of population geneticists to find benefits of recombination sufficient to explain the evolution and persistence of sexual reproduction (Maynard Smith 1971; Barton 2010; Koonin 2016). This nearly unshakable belief arises mainly from the abundant evidence of beneficial transfer events in bacterial genomes. But inferring the value of recombination from successful genomes is like inferring the value of lottery tickets from news about the big winners—the costs to the many losers are hidden from our view.

The discoverers of gene transfer agent chose this name because “… in the absence of evidence for viral involvement we favor considering this system as a bacterial mechanism that exists because of the selective advantage which the capability for genetic exchange might confer.” (Solioz et al. 1975). However, evidence of viral involvement is now abundant (Lang et al. 2017; Shakya et al. 2017), so it is important to reconsider whether GTAs might indeed be just defective prophages, with some functional genes already lost and the rest fated for eventual mutational defects and deletion. Although this could be true for some of the poorly-studied examples of gene transfer agents, it has been ruled out for *α*-proteobacterial GTAs by a detailed investigation of their evolutionary history. Shakya et al. (2017) found that clusters of GTA genes homologous to those in *R. capsulatus* are widespread in a large subclade containing several major orders whose common ancestor is estimated to have lived more than 500 million years ago. D_N_/D_S_ ratios and other features of these genes indicated that they are maintained by the same kinds of selective forces that act on typical chromosomal genes. Similar conclusions were reached by Stanton et al. 2009, Motro et al. 2009, and Guy et al. 2013 about the selective forces acting respectively on *Brachyspira* and *Bartonella* GTA genes. GTAs clearly arise by deletion and recombination processes that place prophage structural and DNA-packaging genes under the control of cellular regulators, and these results suggest they are actively maintained by natural selection acting on benefits they confer.

The modeling we have done shows that lysis of GTA-producer cells imposes a selective cost that overwhelms potential benefits of genetic exchange, but how solid is the evidence that lysis is needed for GTA release? First, all known dsDNA phages lyse their host cells to release their phage; to do this they encode endolysis and holin proteins that disrupt the host cell envelope (Young et al. 2000). Proteins with strong similarity to these are required for RcGTA release; they are encoded separately from the main GTA cluster but induced by the same *cckA* regulation (Westbye et al. 2013). Consistent with a cell lysis function of the endolysin, overexpression in *E. coli* causes cell lysis, and a knockout decreases the release of cytoplasmic components into the medium during GTA production. In *Bartonella*, cells that express the GTA genes are usually unable to form a colony, suggesting that their viability is severely compromised (Québatte et al. 2017).

The cost of cell lysis is only one of the factors that prevent GTA gene clusters from ‘selfishly’ maintaining themselves by transfer into GTA^−^ cells. Another obvious obstacle is the small capacity of most GTA capsids relative to the sizes of the gene clusters encoding them. For most of the known GTAs the capsid capacity is less than half of the gene cluster size (summarized in Lang et al. 2012); the only exception is BaGTA, whose 14 kb cluster length just matches its head capacity (Berglund et al. 2009; Guy et al. 2013). A related problem is that, in systems where this has been investigated, GTA clusters do not contain all the genes needed for GTA production. The RcGTA genes for head spikes, tail fibers, and the holin/endolysin machinery are all encoded separately from its main cluster (Hynes et al. 2012; Brimacombe et al. 2013; Westbye et al. 2016), and the *Brachyspira hyodysenterarae* tail genes are separated from its main cluster (Stanton et al. 2009).

In principle, preferential replication of the GTA gene cluster or preferential packaging of GTA genes into particles could help overcome the cost of lysis. However, the GTA gene clusters lack the replication components of typical prophages, so they are not over-represented in the DNA available for packaging. Those GTA clusters that have been investigated also lack the equivalent of a *cos* DNA-packaging site and a *cos*-recognition protein, which would be needed for preferential packaging. In fact, when the distribution of chromosomal genes in GTA particles has been measured, the RcGTA genes have been found to be under-represented, by about 25% in RcGTA particles (Hynes et al. 2012), and the *Dinoroseobacter shibae* GTA genes by about 75% in DsGTA particles (Tomasch et al. 2018).

### Benefits of general recombination

Our simulations of mutation accumulation in GTA^+^ populations identified two factors capable of overcoming the cell lysis cost of GTA production: synergistic epistasis between mutations and high-fitness regulation of GTA production and recombination. However, the strong synergistic epistasis needed to generate substantial population-level benefits of recombination has not been seen in real bacteria. Bacterial mutation-accumulation experiments have shown that interactions between real genes do not typically generate synergistic epistasis (Elena and Lenski 1997; Maisnier-Patin et al. 2005; Perfeito et al. 2014; D’Souza et al. 2015). Instead, the antagonistic and synergistic epistatic effects seen for different genes largely cancel out, so the net effect is little or no epistasis.

Our results showed that substantial recombination benefits can be also generated by regulation that restricts GTA production and recombination to the most-fit members of the population. Although such regulation is easy to simulate, it would not be easy to implement using the known bacterial regulatory systems, which are adapted to detect specific directly-remediable problems and which. even in combination, would not detect most of the declines in fitness caused by random mutations.

Human brains are proficient at developing ‘adaptive just-so stories’ to explain the phenotypes they observe (Kipling 1902). For *R. capsulatus*, regulation by two-component systems and quorum sensing has been suggested to increase the efficiency of GTA transfer under conditions of reduced fitness (Westbye et al. 2017). This in turn would “increase gene flow in a large community of closely related cells under stressful conditions, increasing the likelihood that allele combinations resulting in favourable traits are combined in one genome.” The opposite argument has been made for Bartonella, with regulation by the ‘alarmone’ ppGpp and by cell division said to be “tuned toward achieving high rates of DNA exchange between the fittest bacteria within a population” (Québatte et al. 2017). Despite these very different regulators and rationales, the kinetics of RcGTA and BaGTA production in lab cultures are strikingly similar, with a strong pulse of GTA release while cells are still in exponential growth and a weaker pulse after growth has ceased (compare Fig. 4 of Solioz and Marrs 1977 and Fig. 1B of Québatte et al. 2017). The identification of very different regulators by molecular investigations may in part be because activators have been sought for *R. capsulatus* and repressors for *Bartonella*. The ppGpp alarmone that regulates Bartonella GTA production (Québatte et al. 2017) might have evolved to prevent GTA production by cells carrying mutations that affect amino acid supply, but it might also simply ensure that the cells’ amino acid supply will be sufficient for synthesis of the capsid proteins.

This work was unable to investigate a third way that benefits of recombination can arise – compensating for stochastic loss of functional alleles in small populations (Muller’s Ratchet, Muller 1964). In small populations of bacteria, horizontal gene transfer can slow the accumulation of harmful mutations (Takeuchi et al. 2014). However, the magnitude of the effect is similar to that of the processes we investigated in large populations, and thus is no more likely to overcome the cost of lysis than are benefits arising from epistasis or regulation.

Although model benefits generated by appropriate epistasis or regulation could overcome the lysis cost of GTA production, they depended on highly optimistic assumptions about the efficiencies of GTA production and recombination. Many factors are likely to reduce real efficiencies far below the 20% limit we found in simulations that incorporated both type of benefits. First, only a fraction of a producer cell’s genome is likely to be packaged. A burst size of ~100 GTA particles per producer cell (typical of the DNA-phage relatives of GTAs) would leave about 90% of the genome unpackaged. Packaging an entire genome into GTA particles would require an unusually efficient packaging mechanism and production of proportionately more head and tail proteins. Although replicating the cell’s genome before packaging would increase the potential burst size, it would also increase the demands on the GTA-production machinery. Second, most particles may not survive long enough to attach to a recipient cell - the discoverers of RcGTA reported that the particles were destabilized by what were thought to be innocuous variations in culture and storage conditions (Solioz et al. 1975). GTA particles will also be lost to diffusion; unless suitable recipients are quite dense many particles may never encounter a recipient to attach to. Dense cultures may create a different problem if recipients are so closely related that transferred fragments rarely carry a different genotype, and quorum-sensing regulation may act to optimize these competing density effects. Another inefficiency arises because the uptake mechanism allows only one of the two DNA strands in the GTA particle to enter the cytoplasm and recombine with the recipient chromosome. This difficulty arises because entry of GTA-borne DNA into the recipient cytoplasm requires the competence gene homologs *comEC* and *comF*, which transport only a single DNA strand into the cytoplasm (the other strand is degraded) (Brimacombe et al. 2015; Québatte et al. 2017). The resulting strand-displacement recombination will thus only change the genotype of one daughter cell, reducing the maximum recombination efficiency to 50%. Recombination efficiency will be further reduced by the leading-end strand degradation that occurs when the competence machinery transports DNA (Pifer and Smith 1985).

The availability of data from co-culture experiments can be used to estimate real transduction efficiencies under laboratory conditions. Marrs and coworkers obtained 2.8 × 10^-5^ doubly-resistant recombinants when they co-cultured *R. capsulatus* cells carrying point mutations giving resistance to either streptomycin or gentamycin (Marrs 1974). Assume that 0.01 of the culture produced RcGTA particles (Fogg et al. 2012)]. If efficiency was perfect, 5 × 10^-3^ of the culture would then have become doubly resistant (see explanatory diagram in Supp. Fig. F). The observed double-resistance frequency was only 2.8 × 10^-5^, so the efficiency must have only been about 5.6 × 10^-3^ transductants per producer cell. If we instead assume that only 0.1% of the cells had produced GTA we can raise the estimated efficiency tenfold. In similar experiments, Quebatte and coworkers obtained 1.0 × 10^-4^ doubly-resistant recombinants when they mixed equal number of *Bartonella* cells carrying a Km^R^ cassette and a Gm^R^ cassette (Québatte et al. 2017). They reported that 6% of the cells had produced GTA, which gives a calculated efficiency of about 3 × 10^-3^ transductants per producer cell. Such low efficiencies could not maintain GTA genes by their recombination benefits even with the most favourable epistasis and regulation conditions.

McDaniels *et al*. reported extraordinarily high transduction frequencies in natural populations of marine bacteria, using GTAs produced by lab cultures of *Roseovarius nubinhibens* and *Reugeria mobilis* (McDaniel et al. 2010). However the reported multiplicities of infection (0.1-10 GTA particles per cell) and frequencies of transduction by a kanR marker (2.4× 10^-2^ – 4.7× 10^-1^) imply transduction efficiencies greater than 100%, which suggests that processes other than GTA transduction may be responsible.

The gold standard for showing that a specific allele or other genetic variant is evolutionarily favoured is analysis of invasion and stability conditions. Tests of invasion determine whether individuals with the variant can increase in frequency after being introduced into a large population of individuals with the standard genotype, and tests of stability determine the converse, whether individuals with the standard genotype can increase in frequency when introduced into a large population with the variant genotype. The analyses in Fig. 9 show that GTA^+^ cells failed both tests when efficiency was as high as 20%, and succeeded at higher efficiencies only when they could sequester the benefits of GTA particles away from the GTA^−^ cells.

Our model was designed to have broad applicability, but some features of the *Bartonella* GTA system deserve special consideration. Although BaGTA, like RcGTA, randomly packages the available genomic DNA, runoff DNA replication from a phage origin of replication 20 kb from the GTA gene cluster causes the GTA genes and adjacent DNA to be overrepresented in GTA particles (~2-fold to 8-fold). The ~2-fold overrepresentation of the GTA genes is not sufficient to allow their selfish maintenance, but the adjacent region contains genes important for host interactions, and it has been suggested that the runoff replication has evolved to enhance the benefits of exchanging these genes (Berglund et al. 2009). However, the conclusions of our recombination model still apply in this situation, especially since *Bartonella* spp. are facultative intracellular pathogens with limited opportunities for DNA transfer.

What forces maintain the gene transfer agent genes in real bacteria, if the benefits of gene transfer cannot do it? We do not know. The simplest explanation would be a direct benefit of the GTA genes to the cells carrying them. However, such a benefit would need to act when the genes are not fully activated, and to be strong enough to compensate for the cost of lysis when they are. It would also need to account for maintenance of all the genes needed for GTA production, so benefits arising from co-option of a single phage-derived gene would not suffice.

Might GTA particles benefit recipients in ways that do not depend on general recombination? One attractive possibility is that GTA particles may serve as a vaccine system when a population first encounters a new phage. If infected cells packaged fragments of phage DNA into GTA particles before lysing, those particles might transmit the phage DNA into uninfected cells, where it could trigger development of CRISPR-cas9 immunity.

Modeling of evolutionary processes is a powerful tool for constraining hypotheses about the consequences of genetic processes and natural selection, but it can never replace either laboratory experiments or studies of natural populations. Our model showed that efficiency of transmission is a critical factor, so we now need experimental data on lysis effects, burst sizes, particle stability and recombination efficiencies. The detailed phylogenetic studies of the RcGTA and BaGTA gene families have shown that GTA genes evolve under the same kinds of constraints as other chromosomal genes(Shakya et al. 2017; Tamarit et al. 2018), but their actual contributions to cellular fitness can only be determined at the laboratory bench.

## METHODS

The R scripts used for all simulations are available in the GTA Evolution project on the Open Science Framework: https://osf.io/krs7h/files/. All simulations except those involving competition between GTA^+^ and GTA^−^ cells used the R script **GTA.R**. Invasion of a large GTA^+^ population by GTA^−^ cells used **GTA-stability.R**, and invasion of a large GTA^−^ population by GTA^+^ cells used **GTA-invasion.R**. Supp. Fig. A shows a flow chart of the GTA.R model. Each run began with a population having no mutations, which accumulated over the generations under the combined effects of new mutation and selection. For runs with no epistasis the maximum number of mutations considered was 35. For runs with epistasis this was increased (antagonistic epistasis) or decreased (synergistic epistasis), to ensure that no more than 10^-9^ of the population was lost to truncation at each generation. A DNA matrix was calculated before each run, to store the probabilities ***P*_*ij*_** that DNA fragments of size ***r*** from cells with ***i*** mutations will contain ***j*** mutations. ***P*** = i!/((j! * (i-j)!) * r^j^ * (1-r)^i-j^ Within each generation the steps were as follows:

### Selection

To simulate selection, the fraction of each mutation class that survives this step is calculated by a fitness function with the form ***w*** = (1 - ***s***)**^*i^α*^**, where ***i*** is the number of mutations in the cell’s genome and **α** is an epistasis coefficient that specifies how interactions between mutations affect fitness. Specific fitness functions used are shown in Fig. 3.

### Production of GTA particles

The regulation function determines the fraction of each mutation class that may produce GTA particles, and pGTA determines the subfractions of these fractions that actually do produce GTAs. The distribution of mutations in the GTA particles is calculated as the product of the DNA matrix and these subfractions. This distribution is then randomized about its mean to simulate uptake of multiple short DNA fragments rather than a single long one of length pGTA.

### Lysis

To simulate the loss by lysis of the cells that produce GTA, the subfraction of cells that produces GTA particles is subtracted from each mutation class.

### Recombination

Simulating recombination requires two steps at each generation. First the DNA matrix is used to calculate, from the GTA distribution, a generation-specific Recombination matrix (Rec_matrix) that gives, for each mutation class, the distribution of changed genotypes after GTA-mediated recombination. Rec_matrix is then applied to the distribution of mutation classes after the lysis step, to calculate the new mutation distribution.

### Mutation

The random distribution of newly arising mutations is calculated using the Poisson distribution. After these are added to the post-recombination distribution, this distribution is used to begin the next generation.

Each run was terminated when the decrease in fitness from one generation to the next was smaller than 10^-9^.

## ACKNOWLEDGEMENTS

We thank Sherwood Casjens for a helpful discussion of capsid sizes and DNA capacities, Olga Zhaxybayeva, Tom Beatty and Andrew Lang for comments on the manuscript, and Sanna Koskiniemi for suggesting the CRISPR-cas9 hypothesis. This work was supported in part by Simons Foundation award #327936 to Olga Zhaxybayeva.

## AUTHOR CONTRIBUTION STATEMENT

RR and SS designed the model. RR did the programming and wrote the first draft of the model. RR and SS polished the manuscript.

